# MicroRNA miR-1002 enhances NMNAT-mediated stress response by modulating alternative splicing

**DOI:** 10.1101/641944

**Authors:** Joun Park, Yi Zhu, Xianzun Tao, Jennifer M. Brazill, Chong Li, Stefan Wuchty, R. Grace Zhai

## Abstract

Understanding endogenous regulation of stress resistance and homeostasis maintenance is critical to developing neuroprotective therapies. Nicotinamide mononucleotide adenylyltransferase (NMNAT) is a conserved essential enzyme that confers extraordinary protection and stress resistance in many neurodegenerative disease models. *Drosophila Nmnat* is alternatively spliced to two mRNA variants, RA and RB. RB translates to protein isoform PD with robust protective activity and is upregulated upon stress to confer enhanced neuroprotection. The mechanisms regulating alternative splicing and stress response of NMNAT remain unclear. We have discovered a *Drosophila* microRNA, *dme-miR-1002*, which promotes the splicing of NMNAT pre-mRNA to RB by disrupting a pre-mRNA stem-loop structure. While NMNAT pre-mRNA is preferentially spliced to RA in basal conditions, miR-1002 enhances NMNAT PD-mediated stress protection by binding via RISC component Argonaute1 to the pre-mRNA, facilitating the splicing switch to RB. These results outline a new process for microRNAs in regulating alternative splicing and modulating stress resistance.

## INTRODUCTION

Neuronal stress resistance is a critical component of health and activity. For many animals, neurons produced during development cannot be replaced and must last for the lifetime of the organism (Asatryan and Bazan, 2017). These neurons withstand a plethora of cellular stressors to maintain viability in the face of disease (Orsini et al., 2016), environmental afflictions (Kagias et al., 2012; Kim and Jin, 2015; Saxena and Caroni, 2011), and aging (Chandra et al., 2017; Chandran et al., 2017; Kirkwood, 2003). Understanding how neurons combat stress has wide-ranging implications for organismal health and quality of life.

Nicotinamide mononucleotide adenylyltransferase (NMNAT) is a housekeeping enzyme long been known as the master enzyme in nicotinamide adenine dinucleotide (NAD^+^) biosynthesis in all living organisms (Zhai et al., 2009). In addition, NMNAT proteins are essential for neuronal maintenance and protection (Ali et al., 2012; Brazill et al., 2017; Fang et al., 2012; Kitaoka et al., 2013; Press and Milbrandt, 2008; Rossi et al., 2018; Sasaki and Milbrandt, 2010; Sasaki et al., 2009; Wen et al., 2011; Yan et al., 2010; Zhai et al., 2006, 2008). NMNAT has been shown to confer robust resistance to various stressors across many species, including hypoxia (Ali et al., 2011; Wen et al., 2013), heat (Ali et al., 2011), oxidative stress (Chiurchiù et al., 2016; Cobley et al., 2018), disease states (Ali et al., 2012; Rossi et al., 2018; Ruan et al., 2015; Zhai et al., 2008), and aging (Liu et al., 2018).

While humans have three nonredundant *Nmnat* genes, *Drosophila melanogaster* has a single *Nmnat* gene (Ali et al., 2013) that is alternatively spliced into two distinct mRNA variants, RA and RB, giving rise to protein isoforms PC and PD, respectively (Ruan et al., 2015). Isoforms PC and PD share similar biochemical properties but have distinct subcellular localizations and divergent neuroprotective functions; isoform PD is cytosolic and is a better neuroprotector than the RA-translated nuclear PC isoform (Ruan et al., 2015). A heat shock response element has been identified in the promoter of *Drosophila Nmnat*, allowing transcriptional upregulation of NMNAT pre-mRNA during stress conditions by stress transcription factor Heat Shock Factor (HSF) (Ali et al., 2011). While NMNAT pre-mRNA is slightly preferentially spliced to RA, it has been shown that the splicing of NMNAT pre-mRNA is shifted to RB in response to stress. Even though NMNAT pre-mRNA transcription is upregulated, only NMNAT splice variant RB is increased in stress conditions. Thus, a post-transcriptional “switch” is predicted to promote the splicing of NMNAT pre-mRNA to RB and to serve as an effective homeostasis strategy to increase the production of the more protective protein PD isoform under stress (Ruan et al., 2015). However, the identity of the “switch” and the mechanism of regulating pre-mRNA alternative splicing under stress remain unknown.

MicroRNAs are short (~22nt) noncoding RNAs involved in mRNA regulation (Carthew et al., 2017; Chawla et al., 2016). MicroRNAs typically bind complementary nucleotide sequences in the 3’ UTR of target mRNAs through the RNA-induced silencing complex (RISC) (O’Brien et al., 2018), and results in the degradation or translational repression of their targets (Gebert and MacRae, 2018). *Drosophila* Argonaute 1, the homolog of mammalian Argonaute 2, is the major RISC component onto which microRNAs are loaded (Yang et al., 2014). While the vast majority of micoRNA-target interactions result in decreased translation of the target, there are some reports of microRNAs upregulating translation of their target mRNAs (Cordes et al., 2009; Ghosh et al., 2008; Jopling et al., 2005; Lu et al., 2010; Mortensen et al., 2011; Ørom et al., 2008; Tsai et al., 2009; Vasudevan et al., 2007), often through binding the 5’UTR (Jopling et al., 2005; Ørom et al., 2008; Tsai et al., 2009). In addition, microRNAs have been implicated in stress response (Edeleva and Shcherbata, 2013; Matsuzaki and Ochiya, 2018; Münzel and Daiber, 2017; Wu et al., 2011) and homeostasis (Bijkerk et al., 2018; Çiçek et al., 2016). For example, miR-221 can protect human neurons from toxicity-induced stress (Oh et al., 2018), several microRNAs have been implicated in fine tuning the redox stress response (Cheng et al., 2013), and miR-393a enhances stress tolerance (Zhao et al., 2018). We hypothesize that microRNAs are excellent candidates for mediating NMNAT splice variant expression under stress, due to their ability to directly influence mRNA levels port-transcriptionally. We thus seek to identify potential microRNAs that are involved in the regulation of NMNAT-mediated stress resistance.

In this study, we identified *dme-miR-1002*, a microRNA with the ability to regulate altering splicing by binding to NMNAT pre-mRNA and upregulate NMNAT mRNA variant RB. Furthermore, we found that *miR-1002* is critical to *Drosophil*a thermotolerance and the stress response of NMNAT RB. Our findings suggest a previously unknown mode of gene expression regulation by non-coding RNAs, and further reveal a new mechanism of maintaining homeostasis and achieving stress resilience.

## RESULTS

### *miR-1002* knockout flies have decreased NMNAT RB and PD expression

*Drosophila melanogaster* has a single *Nmnat* gene producing two mRNA splice variants, RA and RB, which encode protein variants PC and PD, respectively. The pre-mRNA contains seven exons, with exons 1-4 commonly spliced and exons 5-7 alternatively spliced (**Figure 1A**). Since the enzymatic function required for NAD^+^ synthesis is encoded in exons 1-4, both protein isoforms PC and PD have enzymatic activity (Ruan et al., 2015). Exon 7 encodes a nuclear localization motif, therefore isoform PC including exons 6 and 7 is nuclear localized, while isoform PD including exon 5 is localized in the cytoplasm and confers robust neuroprotective function (Ruan et al., 2015).

**Figure 1.**
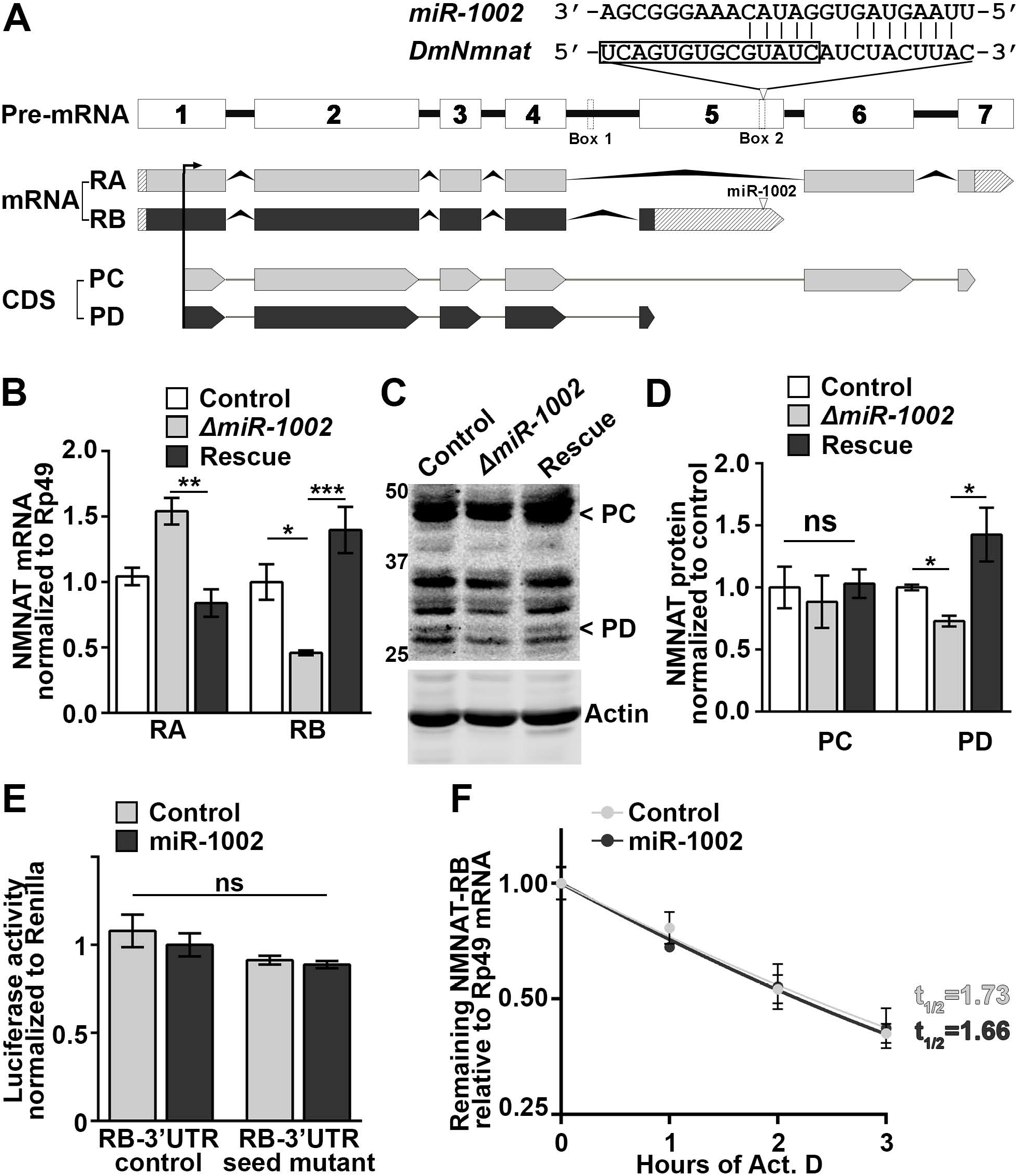
*miR-1002* knockout flies have decreased NMNAT RB and PD expression. (A) Diagram showing *Drosophila* NMNAT pre-mRNA, mRNA splice variants, and protein isoforms. dme-miR-1002 binding sequence on *DmNmnat* depicted at top, with box around Box 2 sequences. (B) NMNAT RNA levels in heads of control wildtype (*yw*), *miR-1002* homozygous knockout (*ΔmiR-1002*), and rescue flies (*ΔmiR-1002; UAS-miR-1002/elav-Geneswitch)*. Geneswitch flies were collected at 2DAE and fed RU486 for 5 days. RP49 used as reference gene. Normalized to control NMNAT RA. Bars indicate mean ± S.E.M. N=4, One-Way ANOVA, Tukey’s post hoc. *p<0.05, **p<0.01, ***p<0.001 (C) Western blot of control, *ΔmiR-1002,* and rescue fly heads. NMNAT PC and PD bands indicated at top. Actin loading control blot at bottom. (D) Quantification of C. Band intensities were normalized to actin, then control NMNAT levels set to 1. Bars indicate mean ± S.E.M. N=4, *p<0.05, One-Way ANOVA, Tukey’s post hoc. (E) S2 cell dual luciferase activity assay. S2 cells were transfected with constructs expressing firefly luciferase linked to control wt NMNAT RB 3’UTR (RB 3’UTR control), firefly luciferase linked to NMNAT RB 3’UTR with the miR-1002 binding sequence mutated (RB 3’UTR seed mutant), or firefly luciferase only. In addition, miR-1002 or a scrambled miR-1002 control plasmid was transfected along with a Renilla luciferase plasmid. Firefly luciferase activity was normalized to Renilla luciferase activity levels, then standardized to the firefly luciferase only condition. Control + wt NMNAT RB 3’UTR was set to 1. Bars indicate mean ± S.E.M. N=4, One-way ANOVA, Tukey’s post hoc. (F) Actinomycin D chase assay. S2 cells were transfected with plasmids expressing miR-1002 or a scrambled miR-1002 control. RNA was collected at indicated timepoints after actinomycin D treatment, then NMNAT RB levels quantified by qPCR. Nonlinear regression of NMNAT RB remaining (normalized to RP49) from initial amount at each time point. Points represent mean ± S.E.M. of N=3 per condition per timepoint. *See also Figure S1.*

To identify potential regulators of *Drosophila Nmnat* expression, we performed a putative microRNA seed sequence scan on NMNAT RB 3’UTR via microRNA.org (Enright et al., 2003; John et al., 2004) and identified *dme-miR-1002* among the highest scoring microRNAs (**Figure 1A**), with a miRSVR score of −0.5339, where miRSVR scores below −0.1 indicate high sensitivity and specificity (Betel et al., 2010).

To test if miR-1002 can modulate NMNAT RB levels, we obtained *miR-1002* homozygous knockout flies (*ΔmiR-1002*) and determined mRNA levels of variant RA and RB in heads by quantitative real-time PCR (**Figure 1B**). We observed significantly reduced levels of NMNAT RB, and increased RA, when compared to wildtype control (*yw*) flies. Furthermore, we found decreased NMNAT PD protein (**Figure 1C** and **1D**) in the knockout flies. When we rescued these flies by conditional overexpression of miR-1002 in the knockout background (*ΔmiR-1002; elav-Geneswitch/UAS-miR-1002*), we observed a significant increase in NMNAT RB and a decrease in RA compared to knockout (**Figure 1B**). The rescue flies also had a significant increase in NMNAT PD (**Figure 1C-D**). This suggests that miR-1002 is important in regulating the expression of NMNAT RB mRNA and PD protein levels *in vivo*.

### miR-1002 shows no effect in altering the stability of NMNAT RB

Reduced variant RB expression in *ΔmiR-1002* suggests that miR-1002 functions to increase the expression of its target mRNA, contrary to conventional 3’UTR-mediated mRNA degradation. To elucidate the molecular mechanisms of transcriptional regulation, we first tested the role of miR-1002 in 3’UTR-mediated mRNA stability using a dual luciferase activity assay (**Figure 1E**). Either the wild-type NMNAT RB 3’UTR sequence (RB 3’UTR control) or NMNAT RB 3’UTR with the miR-1002 seed sequence mutated (RB 3’UTR seed mutant) was inserted downstream of firefly luciferase in an expression vector. This plasmid was co-transfected into *Drosophila* S2 cells along with 2 additional plasmids: a Renilla luciferase expression plasmid (for transfection efficiency control), and either a plasmid overexpressing miR-1002 (miR-1002) or a control plasmid expressing a scrambled miR-1002 sequence (control). 48 hours after transfection, we measured firefly and Renilla luciferase activity to test if miR-1002 can downregulate expression of firefly luciferase through NMNAT RB 3’UTR, and if its binding sequence must be left intact for this to occur. In this reporter assay, we did not observe a significant difference in luciferase activity in cells co-transfected with the miR-1002 plasmid compared to the control plasmid. The same result was observed with both the wildtype and mutant NMNAT 3’UTR luciferase plasmids. This suggests that miR-1002 does not promote 3’UTR-mediated mRNA degradation.

However, it remains possible that miR-1002 can affect the stability of NMNAT RB indirectly through other regulatory pathways. To examine this possibility, we performed an actinomycin D chase assay where miR-1002 or a control was overexpressed by plasmid transfection in S2 cells and the endogenous NMNAT RB degradation rate was monitored after transcription was inhibited with actinomycin D (Qiu et al., 2016; Tew et al., 2014). As shown in **Figure 1F**, no significant difference in mRNA half-life for NMNAT RB was observed between cells expressing miR-1002 and control. Collectively, these results suggest that miR-1002 does not affect the stability of NMNAT RB in a 3’UTR-dependent or an indirect manner.

### *In-silico* analysis reveals the miR-1002-mediated disruption of the pre-mRNA stem-loop and differential minimum free energy values in a sequence-specific manner

To understand how *miR-1002* affects NMNAT expression, we examined its binding sequence relative to known regulatory elements. Two key elements in intron 4 and exon 5 have previously been found to be critical for alternative splicing (Raker et al., 2009). As shown in **Figure 1A**, complementary sequences in intron 4 (Box 1) and exon 5 (Box 2) must bind to form a pre-mRNA stem-loop to facilitate the skipping of exon 5 and inclusion of exon 6, thus forming the RA mRNA variant. When the stem-loop is not formed, exon 5 is preferentially spliced in, producing the RB variant.

The predicted binding sequence of miR-1002 overlaps Box 2 (**Figure 1A**, boxed nucleotides on DmNMNAT); we therefore hypothesized that miR-1002 binding to NMNAT pre-mRNA may interfere with formation of the stem-loop and shift splicing of NMNAT pre-mRNA towards RB. Furthermore, miR-1002 and NMNAT Box 2 sequences are conserved across *Drosophilae* (**Figure S2**). There is remarkable conservation of the Box 2 sequences on NMNAT which miR-1002 is predicted to bind (**Figure S2A**, red asterisks), and the sequence on miR-1002 that are predicted to bind them (**Figure S2B**, red asterisks). This highlights the evolutionary importance of these sequences in regulating NMNAT expression.

To test this hypothesis, we used an *in-silico* approach to predict the RNA secondary structure of various NMNAT and miR-1002 duplexes. In particular, we determined the RNA secondary structure of wild-type exon 4 and intron 5 of NMNAT using RNAfold of the Vienna RNA package (Lorenz et al., 2011), utilizing default parameter settings. To visualize the corresponding minimum free energy structures, we utilized the FORNA web server (Kerpedjiev et al., 2015). We used this platform to model the structural consequences of miR-1002 binding to NMNAT^4-7^, as well as the effects of various mutations on this interaction. As previously indicated (Raker et al., 2009), wild-type NMNAT^4-7^ RNA sequence forms a stem-loop structure via complementary binding of Box 1 (cyan) and Box 2 (blue), a result that we corroborate with our folding of the secondary structure of wild-type NMNAT exon 4 and intron 5 (**Figure 2A**). Assuming that miR-1002 binds its predicted RNA sequence in Box 2, the ability of the corresponding nucleotides to bind other nucleotides in wild-type exon 4 and intron 5 of NMNAT is restricted. Simulating such an assumption, we constrained binding of the corresponding nucleotides using RNAfold, allowing us to observe a massive disruption of the stem-loop structure (**Figure 2B**). The NMNAT^4-7^ mutations modelled in Figures 2C-E encompass mutations within the miR-1002 binding sequence on NMNAT, precluding miR-1002 binding. When miR-1002’s seed sequence outside of the binding stem is mutated (grey, mutated nucleotides in red), miR-1002 can no longer bind, allowing the stem-loop structure to form (**Figure 2C**). When miR-1002’s binding sequence within Box 2 is mutated instead, miR-1002 binding is again impeded (**Figure 2D**). However, the stem-loop is expected to no longer form as a consequence of the disruption of the complementary sequences between Box 1 and Box 2. Finally, when sequences in Box 1 are mutated to be complementary to the Box 2 mutations, the stem-loop forms again (**Figure 2E**). However, corresponding mutations in the seed sequence prevent miR-1002 from binding.

**Figure 2.**
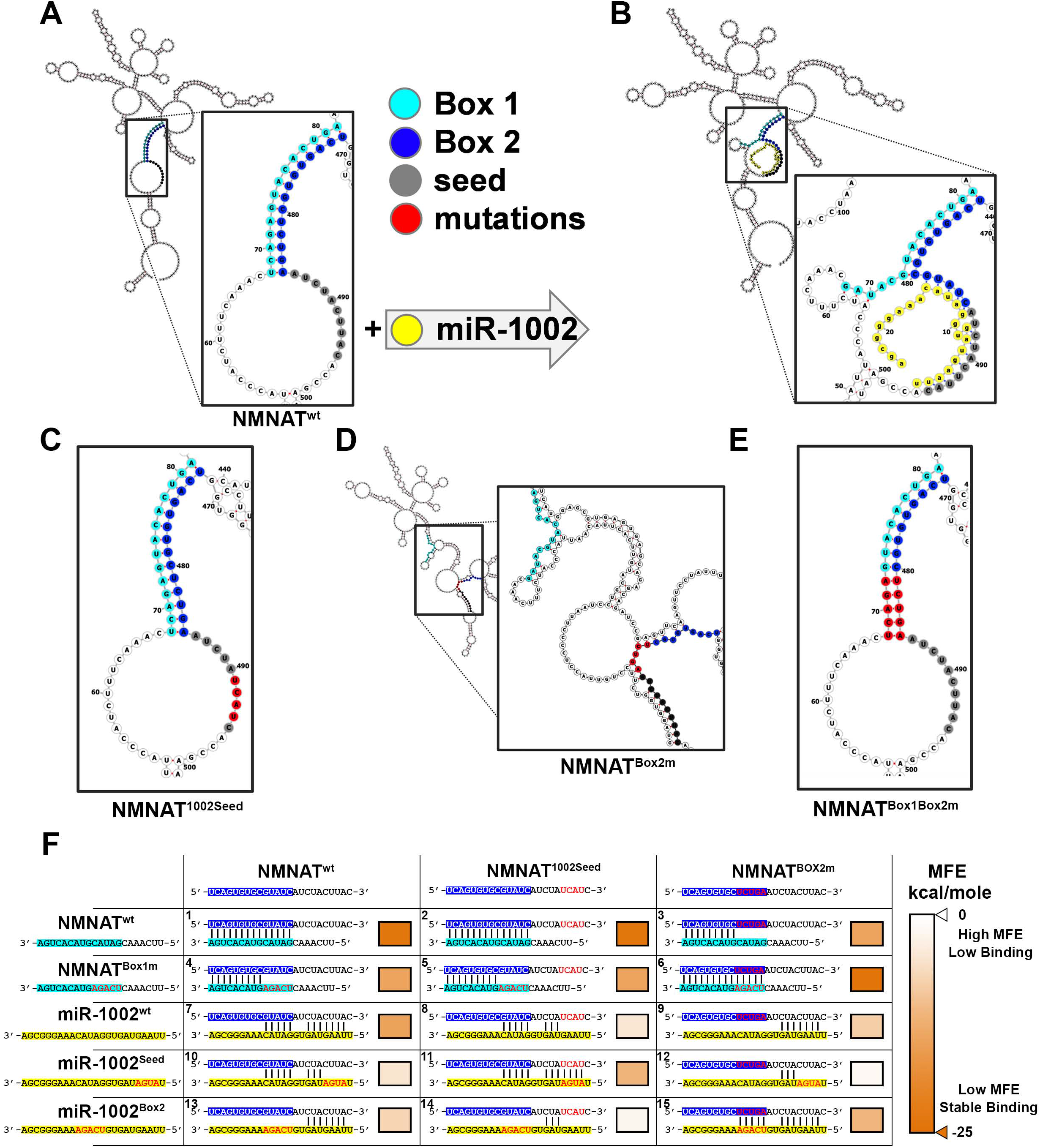
*In silico* analysis reveals miR-1002-mediated disruption of Box 1-Box 2 stem-loop secondary structure and differential MFE values in a sequence-specific manner. (A) FORNA minimum free energy model of the secondary structure of wt exon 4 and intron 5 sequences of NMNAT confirms the emergence of a stem between Box 1 (cyan) and Box 2 (blue) subsequences. (B) miR-1002 (yellow) binds a putative seed sequence (grey) and sequences in Box 2, disrupting the stem. (C-E) FORNA minimum free energy models of NMNAT where red sequences indicated nucleotide scrambled mutations. (C) Scramble mutation to miR-1002 seed (D) scramble mutation to Box 2 (E) Box 1/Box 2 scramble mutation. (F) minimum free energy values (kcal/mole) for various combinations of binding partners for NMNAT Box 2, calculated through duplexfold function in RNAStructure(Reuter and Mathews, 2010). Top row depicts the three NMNAT Box 2 variants used in this analysis-NMNAT^wt^, the wildtype sequence of NMNAT Box 2 region; NMNAT^1002Seed^, NMNAT Box 2 with 4 nucleotides in the miR-1002 seed sequence scrambled; NMNAT^Box2m^, 5 nucleotides in Box 2 were scrambled. The left column depicts the five binding partners used- NMNAT^wt^, the wildtype Box 1 region; NMNAT^Box1m^, 5 nucleotides in Box 1 mutated in a complementary manner to the NMNAT^Box2m^ mutations; miR-1002^wt^, the wildtype miR-1002-5p sequence; miR-1002^Seed^, the seed sequence of miR-1002 mutated in a complementary manner to the NMNAT^1002Seed^ mutations; miR-1002^Box2^, with the Box 2 sequences on miR-1002 mutated in a complementary manner to the NMNAT^Box2m^ mutations. Box to the right of each sequence depicts MFE of duplex binding based on color. *See also Figure S2.*

We then probed the effects of various mutations to NMNAT and miR-1002 on the binding energy of the predicted duplex (**Figure 2F**). Using RNA Structure 6.1 (Reuter and Mathews, 2010), we calculated the minimum free energy of binding for various combinations of binding partners for NMNAT Box 2. The top row depicts the NMNAT exon 5 sequence used in the modelling- wildtype (NMNAT^wt^), seed scramble (NMNAT^1002Seed^, four nucleotides within miR-1002’s seed sequence on NMNAT were scrambled to disrupt miR-1002 binding as in **Figure 2C**), and Box 2 scramble (NMNAT^Box2^, five nucleotides within Box 2 were scrambled to disrupt miR-1002 binding as in **Figure 2D**). The left column shows the binding partners for NMNAT Box 2 used in this model. The top 2 are NMNAT Box 1 wildtype and NMNAT Box 1 scramble (NMNAT^Box1m^, five nucleotides within Box 1 were scrambled in a complementary manner to the Box 2 scramble as in **Figure 2E**). The bottom three are wildtype miR-1002 (miR-1002^wt^), seed scramble (miR-1002^Seed^, four nucleotides within miR-1002’s seed sequence scrambled in a complementary manner to NMNAT^1002seed^, so as to restore miR-1002 binding), and Box 2 scramble (miR-1002^Box2^, five nucleotides within miR-1002’s Box 2 sequence scrambled in a complementary manner to NMNAT^Box2m^, so as to restore miR-1002 binding).

Indeed, when miR-1002’s binding to NMNAT is disrupted through mutations to the binding nucleotides (NMNAT^wt^ + miR-1002^Seed^(10), NMNAT^wt^ + miR-1002^Box2^(13), NMNAT^1002Seed^ + miR-1002^wt^(8), NMNAT^1002Seed^ + miR-1002^Box2^(14), NMNAT^Box2m^ + miR-1002^wt^(9), NMNAT^Box2m^ + miR-1002^wt^(12)), there is a significant increase in minimum free energy (MFE), indicating a less stable structure. MFE decreases when miR-1002 is mutated in a complementary manner to restore binding to NMNAT (NMNAT^1002Seed^ + miR-1002^Seed^(11), NMNAT^Box2m^ + miR-1002^Box2^(15)), indicating a strong and stable structure. In addition, Box 2 forming the stem-loop structure with Box 1 had a low MFE values when the appropriate nucleotides were complementary (NMNAT^wt^ + NMNAT^wt^(1), NMNAT^1002Seed^ + NMNAT^wt^(2), NMNAT^Box2m^ + NMNAT^Box1m^(6)). Together, this analysis demonstrates how alternations in nucleotides have significant effects on RNA secondary structure and binding energy, and that the binding between miR-1002/Box 1 and Box 2 is direct and sequence-specific.

### miR-1002 interferes with the stem-loop mediated alternative splicing of NMNAT pre-mRNA in a sequence-specific manner

To experimentally verify our *in-silico* modelling results in **Figure 2**, we developed a cellular reporter assay to test the functional consequences of various mutations on splicing (**Figure 3A**). *Drosophila* S2 cells were transfected with a reporter plasmid containing firefly luciferase cDNA and NMNAT pre-mRNA exons 4-7 driven from a UAS promoter (pUAS-luc2-NMNAT^4-7^). We co-transfected these cells with a plasmid expressing GAL4 and a scrambled miR-1002 sequence (pAC-GAL4-miR-1002-NegativeScramble) or wildtype miR-1002 (pAC-GAL4-miR-1002). The GAL4/UAS design ensures the co-expression of the Luc-NMNAT^4-7^ reporter and miR-1002-GAL4. We then extracted RNA from these cells and determined splice variant mRNA levels by qPCR. As shown in **Figure 3B**, overexpressing miR-1002 along with wildtype NMNAT reporter (NMNAT^wt^) resulted in a significant increase in RB and decrease in RA (second set of bars) when compared to control (1^st^ set of bars). This is consistent with our hypothesis that miR-1002 interferes with stem-loop formation and shifts the splicing of NMNAT pre-mRNA towards NMNAT RB and away from RA. The upregulation of NMNAT RB by wildtype miR-1002 was not the result of increased transcription, as the RB/RA ratio was significantly increased (**Figure S3A**). When miR-1002 with its seed sequence mutated (3^rd^ set of bars, corresponding to Figure 2F10) or its Box 2 sequence mutated (4^th^ set of bars, corresponding to **Figure 2F13**) was co-transfected along with the wildtype reporter, there was no effect on NMNAT splicing.

**Figure 3.**
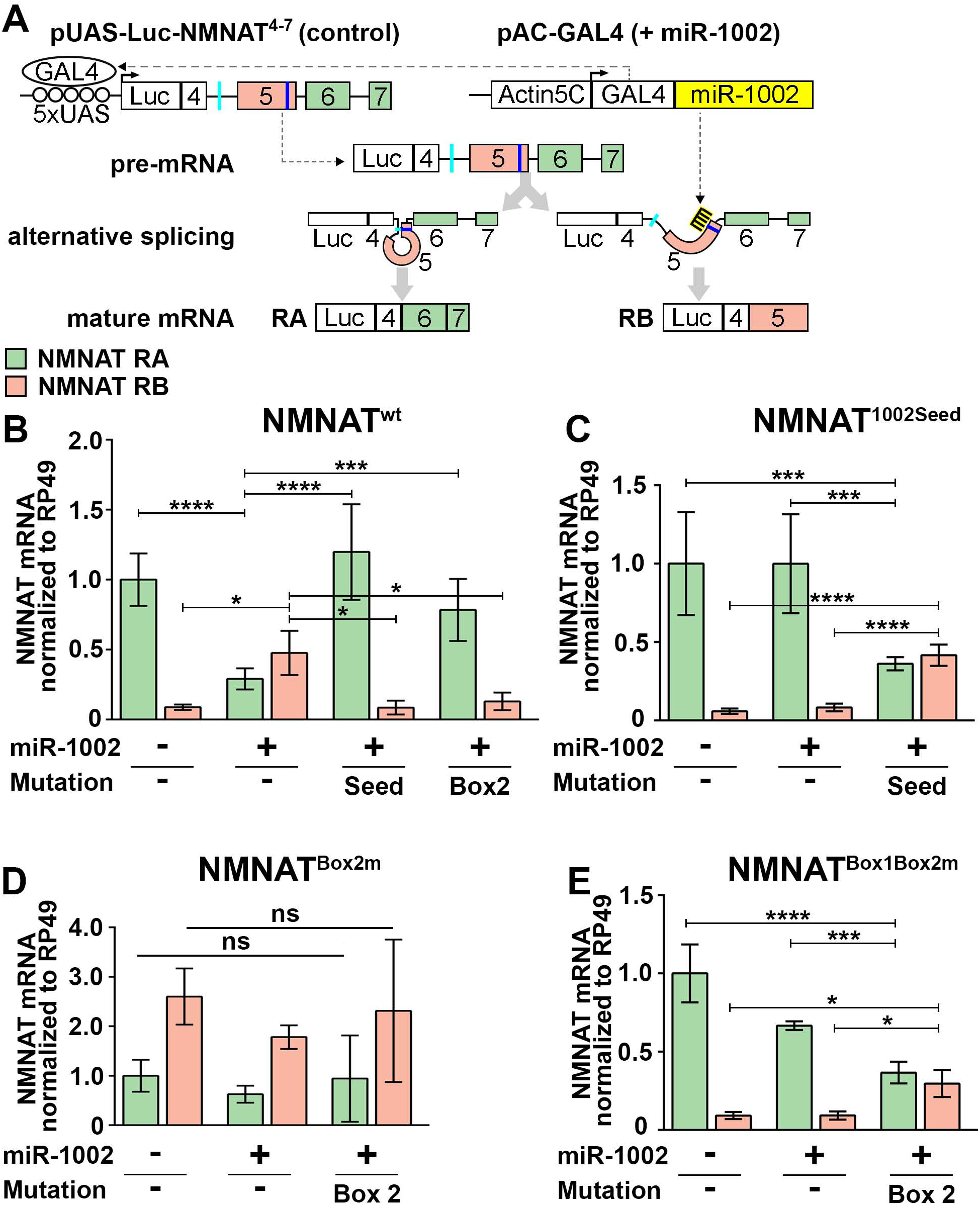
miR-1002 interferes with the stem-loop mediated alternative splicing of NMNAT pre-mRNA in a sequence-specific manner. (A) Schematic representation of S2 cell splicing assay. miR-1002 (yellow) is expressed from a pAC-GAL4 vector. GAL4 protein from pAC-GAL4 activates UAS elements in a reporter construct (pUAS-Luc2-NMNAT^4-7^). Pre-mRNA from the reporter plasmid is then alternatively spliced into 2 variants. RNA is extracted from cells then qPCR is performed for the mature NMNAT mRNA splice variants. This experiment was performed with combinations of the mutations introduced Figure 2. (B-E) Quantification of splicing assay. (B) wt pUASLuc-NMNAT^4-7^ transfected with 3 different PAC-GAL4-miR-1002 variants (miR-1002^wt^, 2^nd^ set of bars; miR-1002^Seed^, 3^rd^ set of bars; miR-1002^Box2^ 4^th^ set of bars) and a negative control miR-1002 scramble plasmid (1^st^ set of bars). (C) pUAS-Luc2-NMNAT^4-7^ Seed scramble (Corresponding to NMNAT^1002Seed^ from Figure 2) transfected with 2 different PAC-GAL4-miR-1002 variants (miR-1002^wt^, 2^nd^ set of bars; miR-1002^seed^, 3^rd^ set of bars) and a negative control miR-1002 scramble plasmid (1^st^ set of bars). (D) pUAS-Luc2-NMNAT^4-7^ Box2 scramble (Corresponding to NMNAT^Box2m^ in Figure 2) transfected with 2 different PAC-GAL4-miR-1002 variants (miR-1002^wt^, 2^nd^ set of bars; miR-1002^Box2^, 3^rd^ set of bars) and a negative control miR-1002 scramble plasmid (1^st^ set of bars). (E) pUAS-Luc2-NMNAT^4-7^ Box1Box2 scramble (Corresponding to NMNAT^Box1Box2m^ in Figure 2) transfected with 2 different PAC-GAL4-miR-1002 variants (miR-1002^wt^, 2^nd^ set of bars; miR-1002^Box2^, 3^rd^ set of bars) and a negative control miR-1002 scramble plasmid (1^st^ set of bars). Bars represent mean ±S.D. transcript levels for each condition and splice variant. Data was normalized to RP49, and control RA set to 1. N>4, One-Way ANOVA, Tukey’s post hoc, *p<0.05, ***p<0.001. ****p<0.0001. *See also Figure S3.*

To determine the sequence-specificity of miR-1002 in regulating NMNAT alternative splicing, we tested the mutations modelled in **Figure 2**. As shown in **Figure 3C**, when the miR-1002 seed sequence outside of NMNAT Box 2 was mutated such that wildtype miR-1002 could no longer bind as in **Figure 2C** (NMNAT^1002Seed^), miR-1002 overexpression had no effect on RB expression (2^nd^ set of bars, corresponding to Figure 2F8). This suggests that miR-1002 binding to its seed sequence is required for its effect on splicing. When miR-1002 itself was mutated in complementary manner to restore binding, we observed a restoration of the downregulation in RA and upregulation in RB (far right bars, corresponding to **Figure 2F11**). Seed-mutated miR-1002 also increased the RB/RA ratio significantly (**Figure S3B**).

In contrast, when miR-1002’s binding sequence within Box 2 was mutated to disrupt both stem-loop formation and miR-1002 binding as in **Figure 2D** (NMNAT^Box2m^), there was no change in NMNAT variant expression in response to miR-1002 overexpression (**Figure 3D**). This was true with both wildtype miR-1002 (second set of bars) and miR-1002 with complementary mutations to NMNAT’s Box 2 mutations in order to restore binding (3^rd^ set of bars, corresponding to **Figure 2F15**). Additionally, the RB levels and RB/RA ratio (**Figure S3C**) for all conditions increased relative to the control wild-type Nmnat^4-7^ plasmid, as the disruption of stem-loop results in exon 5-inclusion to produce RB. This further confirms the validity of our reporter system in assessing stem-loop formation.

Finally, when sequences in Box 1 were mutated in a complementary manner to the Box 2 mutations as in **Figure 2E** (NMNAT^Box1Box2m^), RB levels decreased (**Figure 3E**) and RB/RA ratio decreased (**Figure S3D**). This was expected since the complementary sequences of Box 1 and Box 2 allow the formation of the stem-loop and the exclusion of exon 5 to form RA. While wildtype miR-1002 could not significantly alter NMNAT alternative splicing compared to control due to its inability to bind NMNAT (first and second set of bars, corresponding to **Figure 2F9**), miR-1002 with a complementary Box 2 mutation was able to upregulate RB and downregulate RA (3^rd^ set of bars, corresponding to **Figure 2F15**). With miR-1002 able to bind to NMNAT again, it could compete with Box 1 for binding to Box 2 and increase the RB/RA splicing ratio (**Figure S3D**). Collectively, these results show that miR-1002 binding is necessary and sufficient to interfere with stem-loop formation and shifts the splicing of NMNAT pre-mRNA to NMNAT RB.

### miR-1002 promotes the inclusion of the RB-exclusive 5th exon in an *in vivo* neuronal NMNAT alternative splicing reporter

To assess if miR-1002 regulates the splicing of NMNAT pre-mRNA *in vivo*, we used a previously described alternative splicing reporter (Ruan et al., 2015), where the expression of AcGFP and DsRed were used as a proxy for NMNAT RA and RB, respectively (**Figure 4A**). This is due to an in-frame AcGFP in the 6^th^ exon and DsRed in the 5^th^ exon.

**Figure 4.**
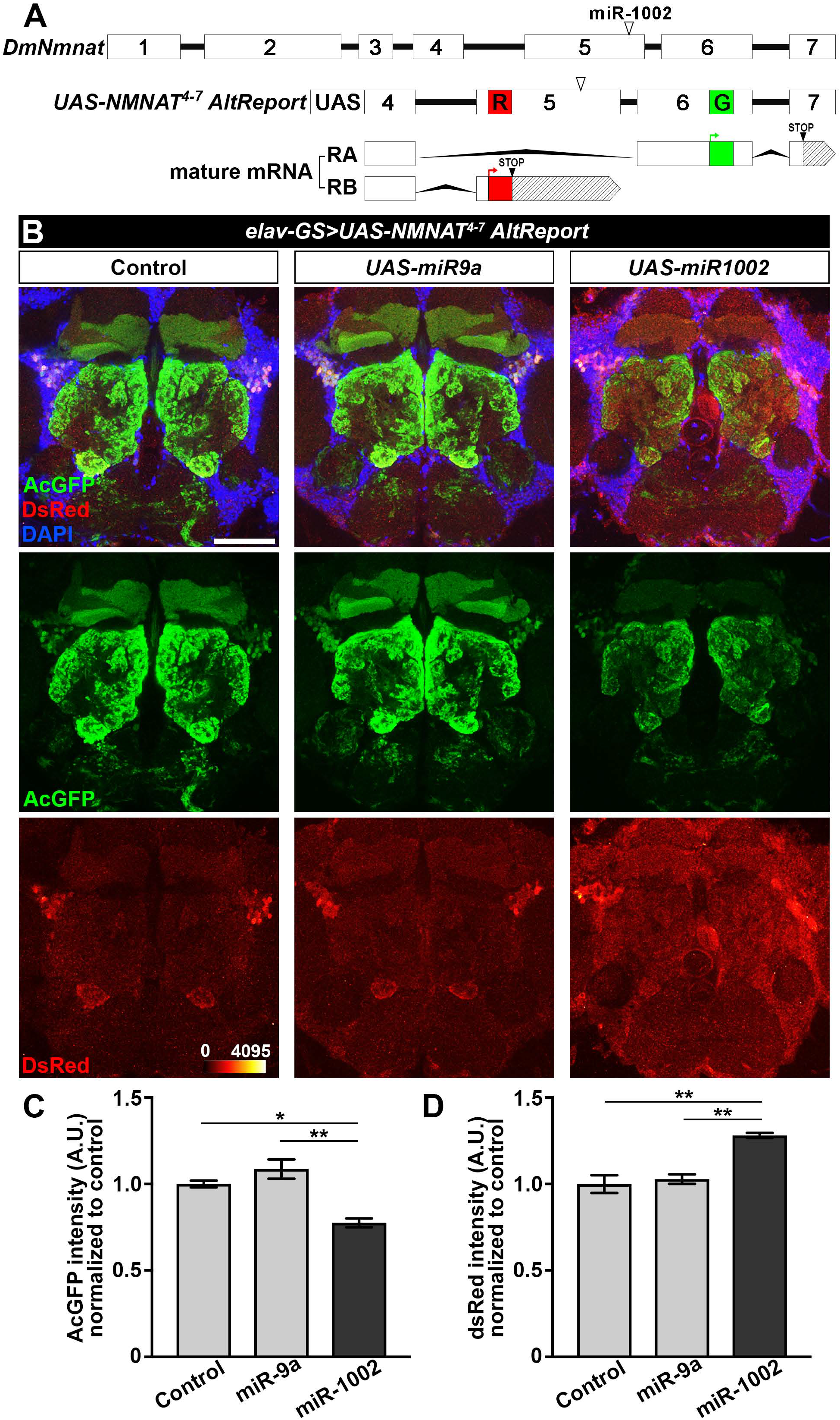
miR-1002 promotes the inclusion of the RB-exclusive 5^th^ exon in an *in vivo* neuronal NMNAT alternative splicing reporter model. (A) Schematic representation of NMNAT reporter (*UAS-NMNAT^4-7^ AltReport*) (Ruan et al., 2015). Genomic *NMNAT* at top with miR-1002 binding sequence indicated with white triangle. *NMNAT alternative splicing reporter* in middle, includes DsRed (Red R) in the 5th exon and AcGFP (Green G) in the 6th exon. Mature mRNA at bottom indicates RA-like splice variant with AcGFP, and RB-like variant with DsRed. (B) Confocal images of fly brains expressing NMNAT reporter and miR-1002, control (Luciferase), or miR-9a. Scale bar 50μm. (C)-(D) Quantification of B. Bars represent mean of 3 midbrain areas per genotype ±S.E.M, normalized to control. N=3, One-way ANOVA, Tukey’s post hoc, *p<0.05, **p<0.01.

Pan-neuronal overexpression of miR-1002 in the brain resulted in early lethality (data not shown). To overcome the lethality, we used an inducible Gene-switch (GS) driver (*elav-GS-GAL4*) to activate expression in adult flies. RU486 (Mifepristone) feeding for 5 days induced expression of the alternative splicing reporter (*UAS-NMNAT^4-7^AltReport*) along with either miR-1002 (*UAS-miR-1002*), a non-targeting control microRNA miR-9a (*UAS-miR-9a*), or negative control luciferase (*UAS-Luc*). AcGFP and DsRed fluorescence in the fly brain was imaged and quantitated. As shown in Figure 4B, we observed a significantly higher DsRed and lower AcGFP fluorescence in the fly brain overexpressing miR-1002 when compared to controls (Figure 4C and 4D), indicating that miR-1002 overexpression promoted the inclusion of the 5th exon of the *NMNAT^4-7^AltReport* splicing reporter construct. These results suggest that miR-1002 overexpression shifts the splicing of NMNAT pre-mRNA towards variant RB in neurons *in vivo*.

### *miR-1002* mediates NMNAT upregulation under stress

NMNAT has been shown to protect many species against a variety of stresses, including heat, hypoxic, oxidative, proteotoxic, and aging stress (Ali et al., 2012; Liu et al., 2018; Miwa et al., 2017; Rossi et al., 2018; Ruan et al., 2015; Zhou et al., 2016). Furthermore, it has previously been reported that *NMNAT* is a stress response gene regulated by the stress transcription factor HSF (Ali et al., 2011). Specifically, neuronal NMNAT PD protein and NMNAT RB mRNA, but not NMNAT RA, are upregulated in response to heat stress, conferring increased self-protection (Ruan et al., 2015). However, the mechanism of the variant-specific stress response “switch” to RB is unknown. miR-1002 presents an excellent candidate for this “switch” since it can bind to NMNAT pre-mRNA to obstruct the stem-loop formation and shift the splicing of NMNAT pre-mRNA into NMNAT RB. In addition, it has been reported that miR-1002 is upregulated in response to heat shock (Funikov et al., 2016) and oxidative stress induced by the treatment of a carcinogenic chemical Cr(VI) (Chandra et al., 2015). We wanted to investigate if the exclusive upregulation of NMNAT RB in these stress conditions is a result of miR-1002 enhancing the splicing of NMNAT pre-mRNA into RB. We subjected wildtype *yw* control and *ΔmiR-1002* flies to heat shock at 37°C for 45 minutes, a stress paradigm that has been shown to induce the highest level of NMNAT RB upregulation (Ruan et al., 2015). As expected, wildtype flies exhibited a significant upregulation in NMNAT RB, but no significant change to RA in response to heat shock, while *ΔmiR -1002* flies lacked the increase in NMNAT RB in response to heat shock (**Figure 5A**). Moreover, there was a significant upregulation in NMNAT RA in the heat shocked *ΔmiR-1002* flies. This supports a model in which NMNAT pre-mRNA transcription is increased during heat stress by heat shock transcription factor HSF, and the pre-mRNA is spliced almost exclusively to NMNAT RB due to increased miR-1002. In the absence of miR-1002, the pre-mRNA is preferentially spliced into RA due to structurally stable (low MFE) stem-loop formation.

**Figure 5.**
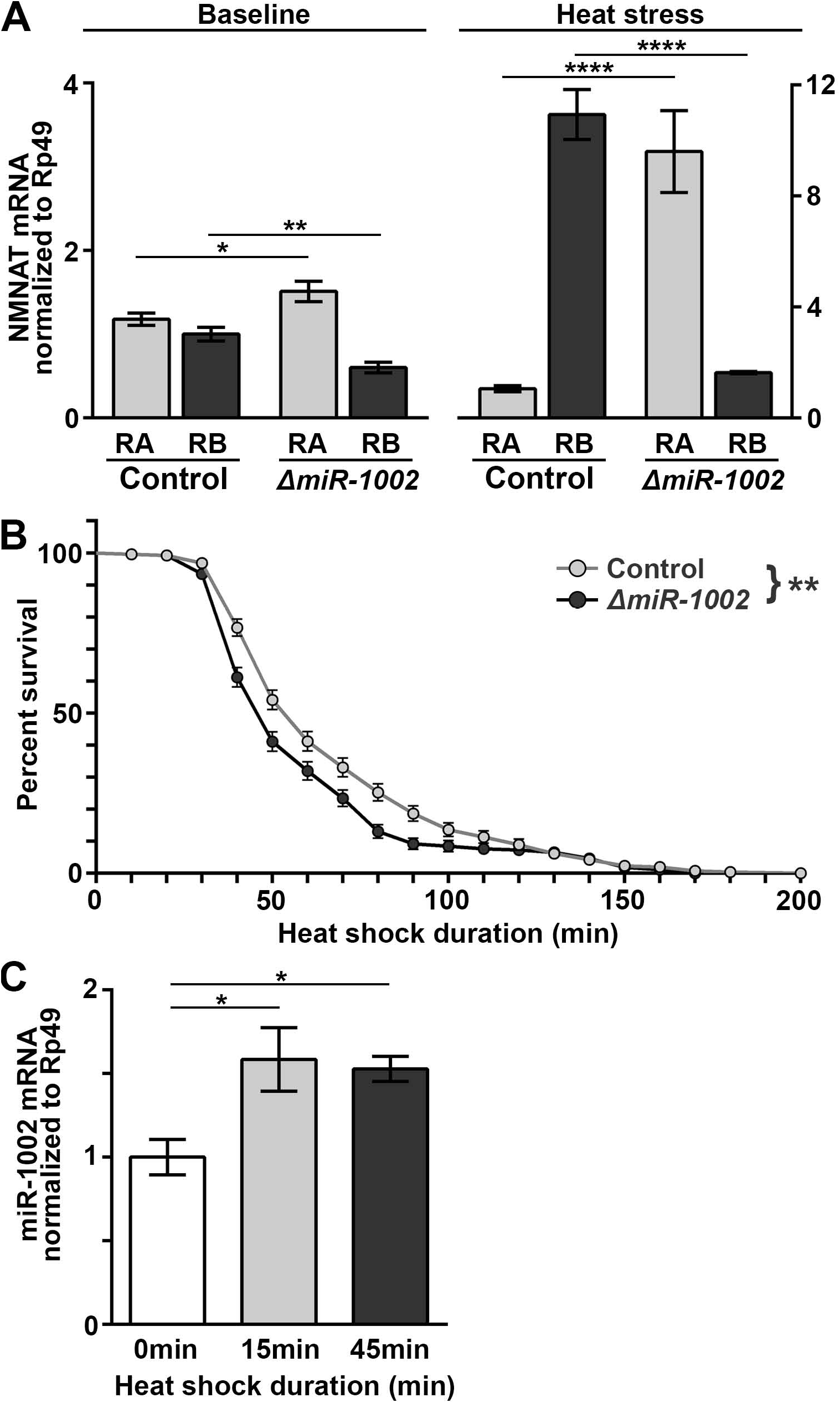
*miR-1002* is critical for thermotolerance and the upregulation of NMNAT RB under heat stress. (A) NMNAT RA and RB levels in control wildtype *yw* and *ΔmiR-1002* flies before and after heat shock. After NMNAT variant expression levels were normalized to RP49, control RB was set to 1. Data is mean of N=7 for baseline, N=4 for heat stress, ±S.E.M. One-way ANOVA, Tukey’s post hoc, *p<0.05, **p<0.01, ****p<0.0001. (B) Thermotolerance assay of control and *ΔmiR-1002* flies. Flies were subjected to heat shock and time taken to paralysis was recorded. Data points represent mean percent active flies at that timepoint. N>200 flies, Log-rank test. p = 0.0031 (C) miR-1002 levels in heat shock. wildtype flies were subjected to heat shock, and miR-1002 levels in heads quantified via qPCR and normalized to RP49 levels. N=4, ±S.E.M. One-way ANOVA, Tukey’s post hoc, *p<0.05.

To assess the functional effects of *miR-1002* in stress response, we performed a thermotolerance assay where flies are subjected to a heat preconditioning (35°C for 30 min) followed by an extreme heat shock at 39°C, and the time to paralysis is recorded. It has been shown that loss of NMNAT leads to reduced latency to paralysis, and that NMNAT overexpression confers paralysis resistance (Ali et al., 2011). As shown in **Figure 5B**, *ΔmiR-1002* flies had a reduced latency to paralysis compared to wildtype flies (p=0.0031). The median survival time for control and *ΔmiR-1002* were 60 minutes and 50 minutes, respectively. At 50 minutes, 54.086 % of control flies remained active, compared to 41.154% of knockout flies. This suggests that miR-1002 is critical for thermotolerance and enhances stress resistance in part though upregulating NMNAT RB as one of the key targets in the stress response network.

In order to determine if miR-1002 is upregulated during heat stress, we extracted small RNA pool from wildtype fly heads after 15 and 45 minutes of heat shock and detected a significant increase in miR-1002 level at both time points of heat shock (**Figure 5C**). This demonstrates that miR-1002 is a heat responsive microRNA, and its increased transcription correlates with increased NMNAT splicing to the RB variant. Higher levels of miR-1002 binding to NMNAT pre-mRNA facilitated efficient splicing to RB under stress.

### Argonaute 1 is essential to miR-1002 regulation of NMNAT splicing

It has been well characterized that during RISC formation, mature *Drosophila* microRNAs are typically loaded onto Argonaute 1 before binding to target transcripts (Förstemann et al., 2007; Pushpavalli et al., 2012). We sought to determine the role of Argonaute 1 in miR-1002-mediated regulation of NMNAT splicing. We obtained Argonaute 1 hemizygous (*AGO1*^+/−^) mutant flies which had previously been shown to have significantly reduced Argonaute 1 levels (Pushpavalli et al., 2014). Heat stress treatment of *AGO1*^+/−^flies resulted in no significant changes to RA or RB levels (**Figure 6A**), suggesting that Argonaute 1 is required to mediate NMNAT stress response.

**Figure 6.**
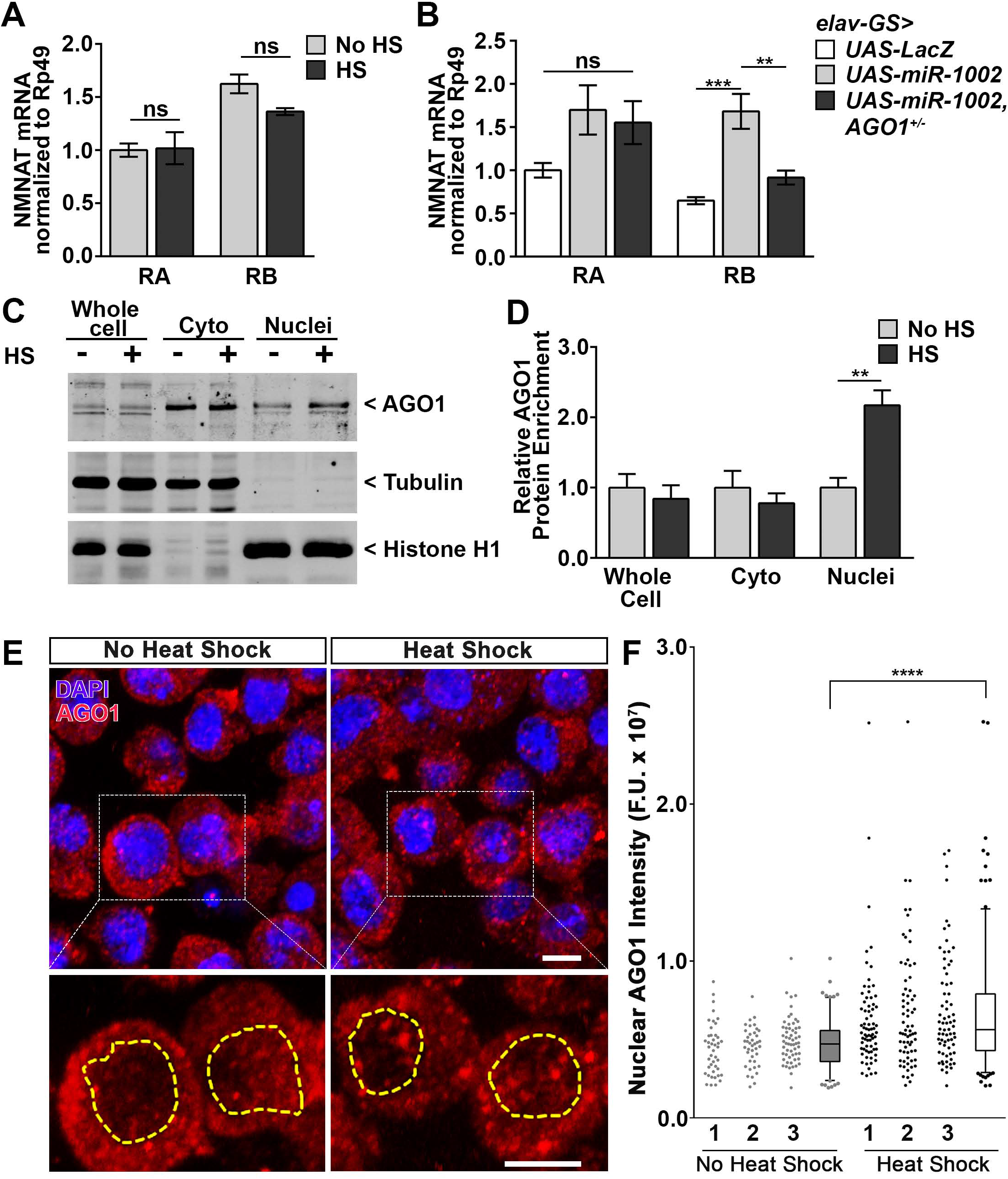
*Argonaute 1* is required for miR-1002-mediated NMNAT regulation. (A) NMNAT splice variants in response to heat shock in AGO1 hemizygous mutant flies. Flies were subjected to 45 minutes of heat shock at 37°C before RNA was extracted, and NMNAT levels normalized to RP49. No HS RA was set to 1. N=4, ±S.E.M. One-way ANOVA, Tukey’s post hoc. (B) NMNAT splice variants of miR-1002 overexpression flies. Control flies expressing LacZ from an elav-geneswitch promoter, flies expressing miR-1002, and flies expressing miR-1002 in an AGO1 hemizygous mutant background were fed RU486 for 5 days, then RNA extracted from heads. NMNAT splice variant levels were normalized to RP49. Control RA was set to 1. N=4, ±S.E.M. One-way ANOVA, Tukey’s post hoc, **p<0.01, ***p<0.001. (C) Western blot of S2 cell fractions before and after heat shock. S2 cells were subjected to 1 hour of heat shock at 37°C, then lysed to purify subcellular fractions. Total cell input, cytoplasm, and nuclei were run together, then blotted for AGO1, Histone H1 (nuclear marker), and α-Tubulin (cytoplasmic marker). (D) Quantification of (C). Band intensities of AGO1 was normalized to Tubulin for the input and cyto fractions, and Histone for the nuclei fractions. Each compartment was normalized to its no HS condition. N=4, ±S.E.M. One-way ANOVA, Tukey’s post hoc, **p<0.01. (E) (F) S2 cells were subjected to 1 hour of heat shock at 37°C, then stained for DAPI and AGO1. Images were taken to measure AGO1 in nuclei. (G) Quantification of (E) and (F). 1-3 are 3 different experiments, with box plot for all experiments combined. Each dot represents a different field. Whispers represent 5 and 95 percentiles. Student’s t-test, ****p<0.0001.

The upregulation of NMNAT RB upon stress can be recapitulated by directly miR-1002 overexpression, as conditional overexpression of miR-1002 in fly brains resulted in a significant increase in NMNAT RB when compared to control overexpressing LacZ (**Figure 6B**). In contrast, when miR-1002 was overexpressed in the *AGO1*^+/−^ background, we observed an abrogation of the NMNAT RB upregulation. This substantiates the essential role of Argonaute 1 in miR-1002 regulation of splicing.

Since most splicing occurs co-transcriptionally in the nucleus (Carrillo Oesterreich et al., 2010), it is likely that AGO1-mediated miR-1002 regulation on splicing occurs in the nucleus. To investigate this possibility, we performed subcellular fractionation of S2 cells before and after heat shock and probed the nuclear, cytoplasmic, and whole cell fractions for Argonaute 1 (**Figure 6C-D**). Argonaute 1 was present in the nuclear fraction (**Figure 6C**), and its level was significantly increased in the nucleus after heat shock (**Figure 6D**). To further validate this finding, we performed immunostaining on S2 cells before and after heat shock. As shown in **Figure 6E-F,** although the majority of Argonaute 1 protein was present in the cytoplasm under normal conditions, nuclear Argoanute 1 was significantly increased after heat shock. Collectively, these results provide evidence for the potential role of Argonaute 1 in the stress-induced nuclear activity of miR-1002 and co-transcriptional splicing of NMNAT.

## DISCUSSION

In this study, we discovered a novel mechanism of a microRNA regulating gene expression under stress. Our results demonstrate that microRNA miR-1002 regulates NMNAT variant-specific expression by interfering with the 4^th^ intron-5^th^ exon stem-loop formation required for pre-mRNA alternative splicing in a direct, sequence-specific manner, rather than by the canonical 3’UTR-mediated mRNA degradation. Additionally, we found that miR-1002 and RISC component Argonaute 1 are essential for NMNAT-mediated neuronal stress resistance.

### A new role for microRNAs in transcriptional regulation—MIMOSAS

This new mechanism of gene regulation, which we term MIMOSAS (<u>MI</u>croRNA-<u>M</u>ediated <u>O</u>bstruction of <u>S</u>tem-loop <u>A</u>lternative <u>S</u>plicing) (**Figure 7**), presents a potential new regulatory layer through which cells can specifically control the expression of transcript variants. While microRNAs are mostly studied in the context of mature mRNA 3’UTR binding and target transcript downregulation, MIMOSAS predicts the binding of microRNAs to interior intronic or exonic regions in the pre-mRNA of targets and regulating mRNA splicing. The outcome of such microRNA-pre-mRNA duplex binding allows mRNA expression in an efficient and splice variant-specific manner. Importantly, MIMOSAS not only extends the functional repertoire of microRNAs, but also adds an exciting, novel example of long-range splicing and RNA secondary structure regulation (Gueroussov et al., 2017; Raker et al., 2009).

**Figure 7.**
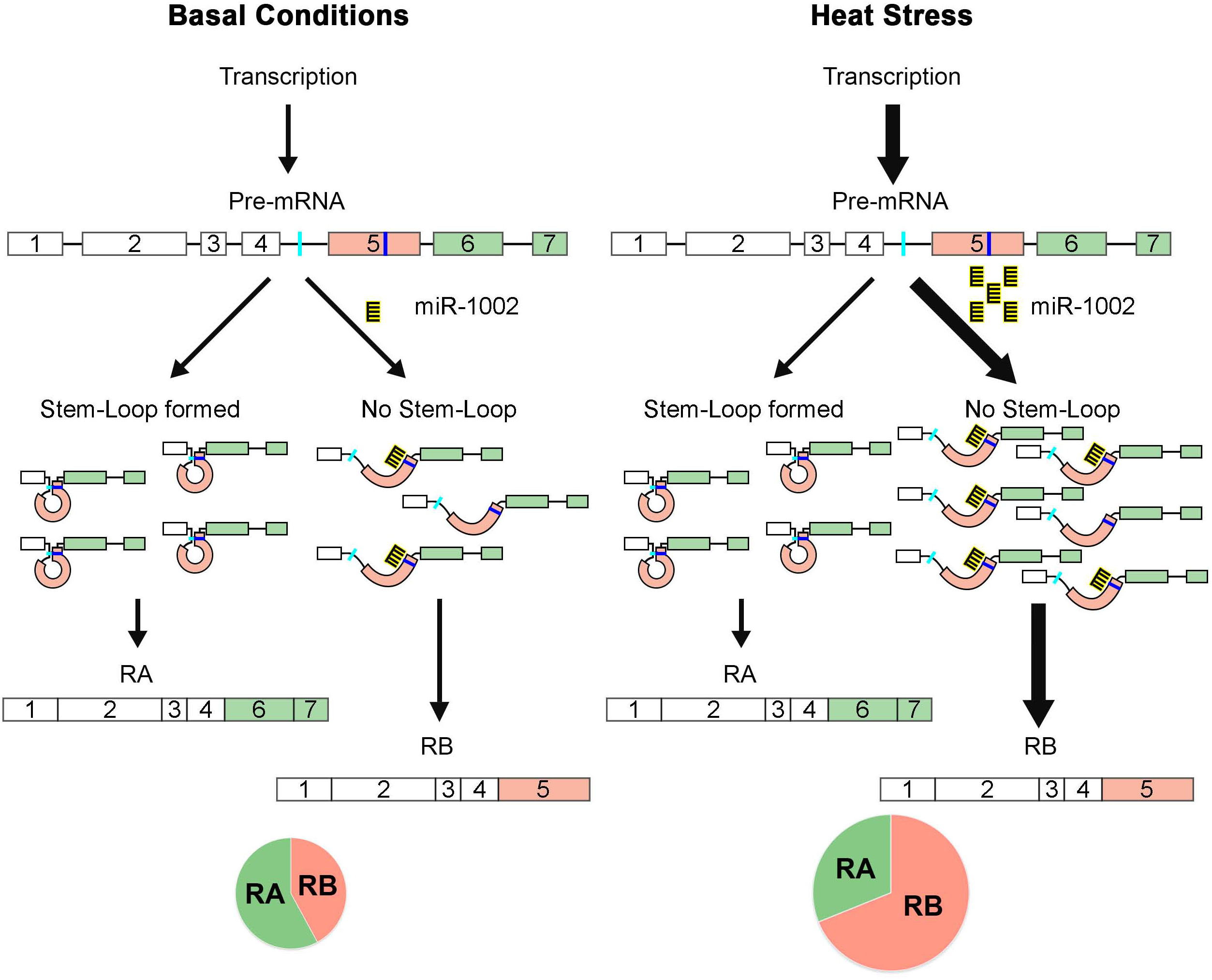
MIMOSAS, a model for miR-1002 regulation of NMNAT splicing. Under basal conditions (Left), NMNAT is preferentially spliced into the RA mRNA variant through stem-loop formation. Under stress (Right), HSF upregulates transcription of NMNAT pre-mRNA. miR-1002 is also upregulated under stress, and miR-1002 enhances the splicing of NMNAT into the RB variant by preventing formation of the stem-loop. This upregulation serves as a stress induced “switch” to increase protection through NMNAT PD.

Previous studies have suggested that non-coding RNAs can affect splicing. Gonzalez, et. al. showed that a lncRNA can promote the retention of a particular exon in FGFR2 alternative splicing (Gonzalez et al., 2015). Double stranded RNAs can affect the splicing of genes associated with muscular dystrophy (Liu et al., 2012). A small nuclear RNA has been found to regulate the splicing of Serotonin Receptor IIC, which is associated with Prader-Willi Syndrome (Kishore and Stamm, 2006).

miR-1002’s ability to regulate splicing of a target transcript offers new insight to how microRNAs control gene expression. It remains to be seen how prevalent this mechanism is, and whether it is conserved. It was previously reported that 42.6% of microRNA target sites in human T-REx 293 cells were found in mRNA CDS regions (Helwak et al., 2013). A study mapping Kaposi's sarcoma-associated herpesvirus (KSHV) microRNA binding sites in B cell lymphoma-derived cells and epithelial cells found that a majority of microRNA target sites were found within the CDS of mRNAs, not the 3’UTR (Gay et al., 2018). Given that approximately 95% of multi-exonic human genes are alternatively spliced (Kim et al., 2018), microRNA target sites in the exonic and intronic regions present ample opportunities for microRNAs to target and influence pre-mRNA splicing.

Our findings suggest that MIMOSAS through miR-1002 is mediated by Argonaute 1. Whether miR-1002 regulation of NMNAT occurs in the nucleus, cytoplasm, or both is important for understanding the mechanism of microRNA-mediated splicing regulation. Since most splicing occurs co-transcriptionally (Oesterreich et al., 2016), the nucleus is the most likely location for such microRNA-mediated regulation. Supporting this hypothesis is the detection of microRNAs in the nucleus. RNALocate lists over 200 examples of experimentally determined nuclear non-coding RNAs in a neuronal or neuron-like cells (Zhang et al., 2017). On the other hand, Argonautes have also been shown to be localized to the nucleus and bind chromatin (Cernilogar et al., 2011; Huang and Li, 2014; Huang et al., 2013; Pushpavalli et al., 2012; Taliaferro et al., 2013). For example, Argonaute has been found to be nuclear localized in many cell types, including T47D (Chu et al., 2010), 10G, ER293 (Ohrt et al., 2008), HEK293, HeLa, and RPE-1 cells (Rüdel et al., 2008). Our study detected Argonaute1 in the nuclear and cytoplasmic fraction, further supporting the notion that microRNA regulation of splicing may occur in both the nuclear and cytoplasmic cellular compartments. Specifically, our observation that the dependence of miR-1002 on Argonaute 1 and the stress-induced nuclear enrichment of Argonaute 1 supports a model in which Argonaute 1-miR-1002 localization to the nucleus during stress conditions facilitates the splicing of NMNAT pre-mRNA into the RB variant under stress.

It is also important to recognize the possibility of cytoplasmic regulation of splicing, given the large body of literature suggesting that splicing events are not exclusive to the nucleus and often occur outside the nucleus, especially in cells with large cytoplasm such as neurons (Back et al., 2006; Bell et al., 2010; Denis et al., 2005; Glanzer et al., 2005; König et al., 2007; Rüegsegger et al., 2001; Yoshida, 2007). For example, it has been reported that 44% of genes expressed in rat dendrites retained introns, 59% of genes in mouse neuronal soma retained introns, 61% of cytoplasmic genes in BAT cells and 49% of genes in cardiomyocytes retained introns (Khaladkar et al., 2013). With cytoplasmic splicing being prevalent, it is possible that miR-1002 regulates NMNAT pre-mRNA splicing in the cytoplasm. Moreover, microRNA loading onto RISC most likely occurs outside of the nucleus. It has been shown that nuclear RISC complexes are deficient in RNA loading (Gagnon et al., 2014), and nuclear activated RISC seems to originate from the cytoplasm (Ohrt et al., 2008). It is likely that microRNA-RISC complex-mediated regulation of splicing happens in multiple steps that occur in multiple subcellular compartments. Such translocation between compartments allows ample opportunity for regulation, while preserving the time efficient response upon stress.

### MicroRNA miR-1002 is a part of the stress response network

Our previous study has found that under stress, NMNAT RB variant is upregulated to produce the neuroprotective protein isoform (PD). Therefore alternative splicing of NMNAT functions as a switch to enhance neuroprotection under stress (Ruan et al., 2015). Excitingly, we discovered microRNA miR-1002 as the main driver of the ‘switch’, as the upregulation of NMNAT RB under stress is eliminated in *ΔmiR-1002* flies (Figure 5A). Specifically, the observation that the heat shock upregulation of NMNAT RA in *ΔmiR-1002* flies (6.23x) mirrored NMNAT RB increase (10.94x) in wildtype flies suggests that miR-1002 efficiently disrupts stem-loop and pushes all newly transcribed pre-mRNA into RB under stress. Without miR-1002, the newly transcribed NMNAT pre-mRNA was almost exclusively spliced into RA.

Our model predicts a novel role for miR-1002’s regulation of NMNAT (**Figure 7**). In basal conditions, NMNAT pre-mRNA is spliced to RA with a slight preference via stem-loop formation (Ruan et al., 2015). Under stress, miR-1002 is upregulated and switches the splicing of newly transcribed NMNAT pre-mRNA almost exclusively to RB. Given the robust neuroprotective function of NMNAT isoform PD, miR-1002-mediated switch of alternative splicing enhances neuronal resistance to stress and confers organismal resilience. This is supported by the reduced thermotolerance observed in *ΔmiR-1002* flies. The modest reduction in thermotolerance suggests that miR-1002 is one of the key players in the complex stress response network. Many other genes and proteins including heat shock proteins (HSPs) have been found to contribute to stress resistance (Bartelt-Kirbach et al., 2016; Jo et al., 2017; Kawasaki et al., 2016; Morrow et al., 2016; Shukla et al., 2014). Importantly, our study found that overexpression of miR-1002 in neurons in the brain was sufficient to shift alternative splicing towards RB *in vivo* further substantiates the role of miR-1002 in enhancing NMNAT-mediated neuroprotection.

Our discovery of a novel mechanism of a microRNA regulating gene expression under stress unveils regulation of gene expression by noncoding RNAs in a state-specific manner and underscores the importance in considering alternative splicing modulation as therapeutic means. As 60% of point mutations associated with diseases have been found to be related to splicing defects (Jensen et al., 2009), several methods have been developed to modify alternative splicing. For example, duplex RNAs have also been used to modify alternative splicing in mammalian cells (Liu et al., 2012), and a method of using single stranded siRNAs has been developed to modify the splicing of a Dystrophin RNA, which is associated with Duchene muscular dystrophy (Liu et al., 2015). The results of our work introduce the possibility of microRNAs as tools to modulating alternative splicing. While more work is required to identify other microRNA and target mRNA pairs, our work shows a promising new direction for conferring neuroprotection and organismal resilience.

## Acknowledgements

We would like to thank Zoraida Diaz-Perez and Kai Ruan for technical support. This research was supported by the Lois Pope LIFE Foundation Fellows Program (J.M.B., Y.Z., C.L.), and National Institutes of Health (NIH) (R56NS095893 to R.G.Z.).

## Materials and Methods

### *Drosophila* Strains and Maintenance

Flies were maintained and crossed in a climate-regulated 25°C incubator in a 12hr light-dark schedule with 60-65% humidity on cornmeal-molasses-yeast medium. *UAS-NMNAT ^4–7^ AltReport* was previously generated in lab (Ruan et al., 2015). *yw* flies were used as wildtype controls throughout this study. Also used were *UAS-miR-1002* (Bloomington stock number 60657), *miR-1002* knockout flies (Bloomington stock number 58949), *UAS-Luciferase* (Bloomington stock number 35788), *UAS-miR-9a* (Bloomington stock number 41138), AGO1 mutant flies (Bloomington stock number 10772), and *elav-geneswitch* (Bloomington stock number 43642).

### Cell Culture

*Drosophila* S2 cells (ATCC) were maintained at room temperature in Schneider’s *Drosophila* medium (Lifetech) supplemented with 10% Fetal Bovine Serum (Atlanta Biologicals) and 1% Penicillin/Streptomycin (GIBCO). S2 cells are derived from embryonic cells and are believed to be male.

### RNA Extraction and qRT-PCR

Female fly heads (30-50 per N) at 2 DAE (days after eclosion) were homogenized using a mechanical pestle homogenizer in 200μl Trizol (Sigma). After adding 800μl Trizol with 200μl chloroform and a 15min centrifugation at 4°C, the RNA layer was collected. Total RNA was extracted using Favorgen RNA mini kit according to the manufacturer’s protocol. Reverse transcription was performed using the Applied Biosystems High Capacity cDNA Reverse Transcription Kit. 2 μg RNA was used in a 20 μl RT reaction according to the manufacturer’s protocol. For qPCR, a 96 well plate was used. 100ng of cDNA from each sample was loaded into each well, with triplicate wells for each biological sample. Custom primers were designed for each target transcript, and Bio-Rad iQ SYBR Green Supermix added according to the manufacturer’s protocol. A CFX connect real-time detection system (Bio-Rad) was used to perform qPCR. The quantification of transcripts was calculated via the 2(-Delta Delta C(T)) method (Livak and Schmittgen, 2001). RP49 was used as reference genes. See table S1 for primer sequences.

### miRNA Extraction and qRT-PCR

30+ female fly heads were collected and homogenized with a mechanical pestle homogenizer in 700 μl Qiazol reagent. After addition of chloroform and 15 min spin at 4°C, the RNA layer was collected. Total RNA including microRNA was extracted using the miRNeasy kit (Qiagen) according to the supplier’s protocol. 200 ng of RNA was used in the RT reaction with the miScript RT kit (Qiagen). qPCR was performed using a custom LNA probe for miR-1002 using the miRCury SYBR green reagent (Qiagen).

### Western Blot

Female fly heads (10 per N) were collected and homogenized in RIPA buffer (Sigma). 4X SDS loading buffer was added, then samples boiled at 95°C for 5 minutes. Samples were loaded onto a 10% SDS gel for electrophoresis. After transfer onto a nitrocellulose membrane (Biorad) and blocking with TBS/Blocking buffer (Rockland), the membrane was probed with anti-DmNMNAT (1:8,000) and anti-β-Actin antibody (1:10,000, Sigma). Membranes were further probed with dye-conjugated secondary antibodies (1:10,000, Rockland), then imaged at IR700 and IR800 on an Odyssey Infrared Imaging system (LI-COR Biosciences). Analysis was performed using Imagestudio (LI-COR).

### Plasmids

pUAS-luc2 was a gift from Liqun Luo (Addgene plasmid # 24343 ; http://n2t.net/addgene:24343 ; RRID:Addgene_24343)(Potter et al., 2010). pAC-GAL4 was a gift from Liqun Luo (Addgene plasmid # 24344; http://n2t.net/addgene:24344 ; RRID:Addgene_24344). pRL-ubi63E was a gift from Alexander Stark (Addgene plasmid # 74280 ; http://n2t.net/addgene:74280 ; RRID:Addgene_74280) (Arnold et al., 2013). *NMNAT RB 3’UTR*, *NMNAT*^*4–7*^, and *miR-1002* were all cloned out of *Drosophila* genomic DNA into PGEM-T-Easy before ligation into the appropriate plasmids. miR-1002 negative scramble was ordered from Genscript for ligation into pAC-GAL4.

Mutations to plasmids were created with Quikchange II site-directed mutagenesis kit (Agilent). Sequences for primers used in mutations are found in table S1.

### Luciferase activity assay

*Drosophila* S2 cells in a 12 well plate were transfected with 3 plasmids using JetPrime- pRL-UBI-63E; pAC-GAL4-miR-1002-negativeScramble, or pAC-GAL4-miR-1002; and pUAS-LUC2-NMNAT-RB-3’UTR, or pUAS-LUC2-NMNAT-RB-3’UTR-miR-1002-seedmut. 48 hours after transfection, cells were collected and lysed with a lysing buffer (100mM KCl, 20mM HEPES, 5% Glycerol, 0.1X Triton-X, 1mM DTT, and Complete protease inhibitor (Roche)). ProMega DUAL-GLO Luciferase assay was performed to measure firefly and Renilla luciferase activity according to the manufacturer’s protocols. Luminescence was measured using a FluoStar Omega microplate reader in a 96 well plate, with triplicate wells per biological sample.

### Actinomycin D chase assay

Actinomycin D (Sigma) was reconstituted in DMSO (Sigma) at a concentration of 1mg/ml. S2 cells were transfected with pAC-GAL4-miR-1002 or pAC-GAL4-miR-1002-scramble. 48 hours after transfection, Actinomycin D was added to the cells at a concentration of 10 μg/ml. Cells were harvested at 0, 1, 2, and 3 hours after actinomycin treatment, then RNA collected for qPCR.

### Minimum Free Energy (MFE) RNA secondary structures

RNA secondary structures were predicted using RNAfold as of the Vienna RNA package. Secondary structures were visualized using the FORNA web server (http://rna.tbi.univie.ac.at/forna/). *Nmnat* exon 4-intron 5 and the various mutations (Figures 3A, 3C-E) were modelled for MFE through Vienna RNA fold (Lorenz et al., 2011). The structure was then visualized using FORNA (Kerpedjiev et al., 2015). miR-1002 was manually inserted in Figure 3B. RNAStructure’s duplexfold function was used to calculate MFE values (Reuter and Mathews, 2010).

### Drug feeding

RU486 (mifepristone, Sigma) was dissolved in ethanol, then mixed into fly food at a concentration of 50μg/ml. Flies expressing an elav geneswitch driver were fed food containing drugs or vehicle control for 5 days before brains were dissected.

### Immunocytochemistry of fly brains

Adult female brains were fixed in 3.5% formaldehyde for 15 min and washed in PBS with 0.4% Triton X-100. Brains were incubated with anti-RFP (Abcam, 1:500) and an alexa-555 conjugated secondary antibody (Invitrogen, 1:250). DAPI (Invitrogen, 1:300) staining was performed post-secondary antibody staining. All antibody incubations were performed at 4 °C overnight in the presence of 5% normal goat serum. Brains were then mounted and imaged on an Olympus IX81 confocal microscope and processed using FluoView 10-ASW (Olympus). Images were quantified using ImageJ. The midbrains of 3 brains per genotype were analysed.

### Stress Paradigms

For heat stress, female flies were collected 2 DAE and subjected to a 15 or 45-minute heat shock in vials placed into a 37°C incubator. Flies were collected after heat shock for RNA extraction from heads.

For thermotolerance assay, 20 flies per vial at 2 DAE were submerged in a 35°C water bath for a 30-minute preconditioning. Flies were immediately moved to a 39°C water bath. Vials were checked under a light microscope every 10 minutes to count paralysed flies.

### Subcellular Fractionation

S2 cells were collected and washed in ice cold PBS. Subcellular fractionation was performed with the NE-PER kit (Thermo-Fisher), according to the manufacturer’s instructions. Both nuclei and cytoplasmic fractions were collected. Western blot was performed for AGO1 (abcam), α-Tubulin (Sigma), and Histone-H1 (active motif).

### S2 cell immunostaining

Coverslips were treated with 20 ug/ml of poly-D-lysine in Hank’s Basic Salt Solution on 35-mm culture dishes overnight. Coverslips were rinsed with S2 cell medium before seeding cells. 24 hours post seeding, heat shock the cells at 37°C for 1 h. Cells were fixed with 4% formaldehyde for 15 min and permeabilized with 0.4% Triton X-100 for 5 min. Coverslips were blocked with 5% goat serum in PBS at 37°C for 30 min and then incubated with AGO1 antibody (1:100) in blocking solution at 37°C for 2 h. After washing with PBS for 5 min, 3 times, coverslips were incubated with Cy5-labeled anti-rabbit secondary antibody (1:500) in blocking solution at 37°C for 1 h. In the second PBS wash, DAPI was added (1:10000), then coverslips mounted with VECTASHIELD Antifade Mounting Medium.

Quantification of nuclear AGO1 was performed as previously described (Brazill et al., 2018). Briefly, images were taken using a 60x oil immersion objective lens. Region of interest (ROI) was selected based on DAPI staining. Auto-segmentation was conducted by interactive h-maxima watershed. Three parameters were adjusted: seed dynamics determined by h-maxima (h=5892), global watershed stop criteria determined by intensity threshold (T=1119), and regional stop criteria determined by peak flooding (%=90). Nuclei were defined as objects larger than 10 μm^2^.

### Quantification and Statistical Analysis

Prism was used for statistical analysis and graph generation. Student’s t-test, one-way ANOVA, and two-way ANOVA with Tukey’s post hoc were used. Log-rank test was used to calculate significant differences between the 2 populations in the thermotolerance assay. All analyses were two-tailed. *p<0.05, **p<0.01, ***p<0.001, ****p<0.0001

### Data and Software Availability

Putative microRNA seed sequence scan was performed using (http://microRNA.org) (Betel et al., 2008, 2010; Enright et al., 2003; John et al., 2004). RNA secondary structures were modelled using RNAfold as of the Vienna RNA package (Lorenz et al., 2011). Secondary structures thus obtained were visualized using web-based FORNA (http://rna.tbi.univie.ac.at/forna/) (Kerpedjiev et al., 2015). RNA duplex minimum free energy values were calculated using RNAStructure (Reuter and Mathews, 2010)

