## Supplementary figures and images for "MicroRNA miR-1002 enhances NMNAT-mediated stress response by modulating alternative splicing"

### Supplemental Figure 1

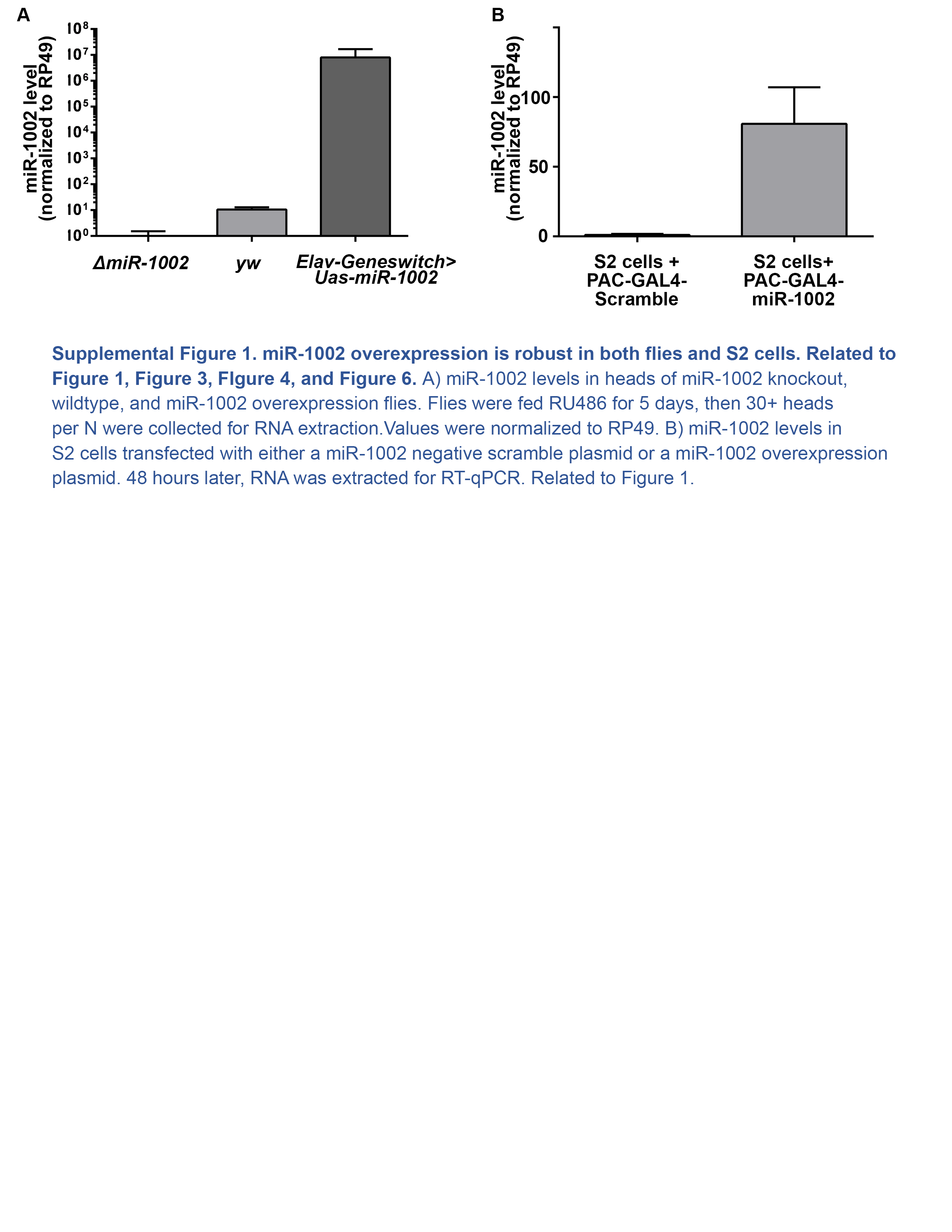

### Supplemental Figure 2

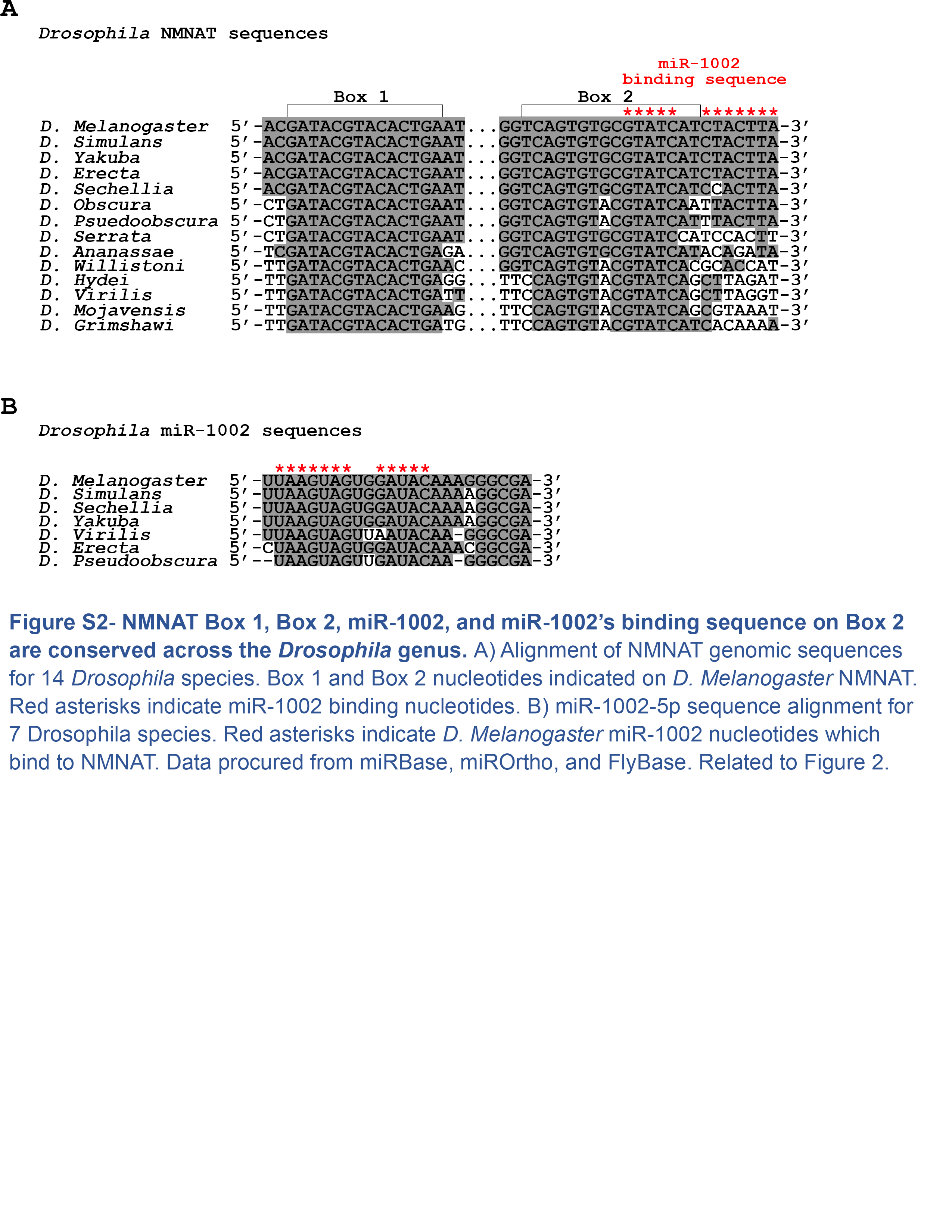

### Supplemental Figure 3

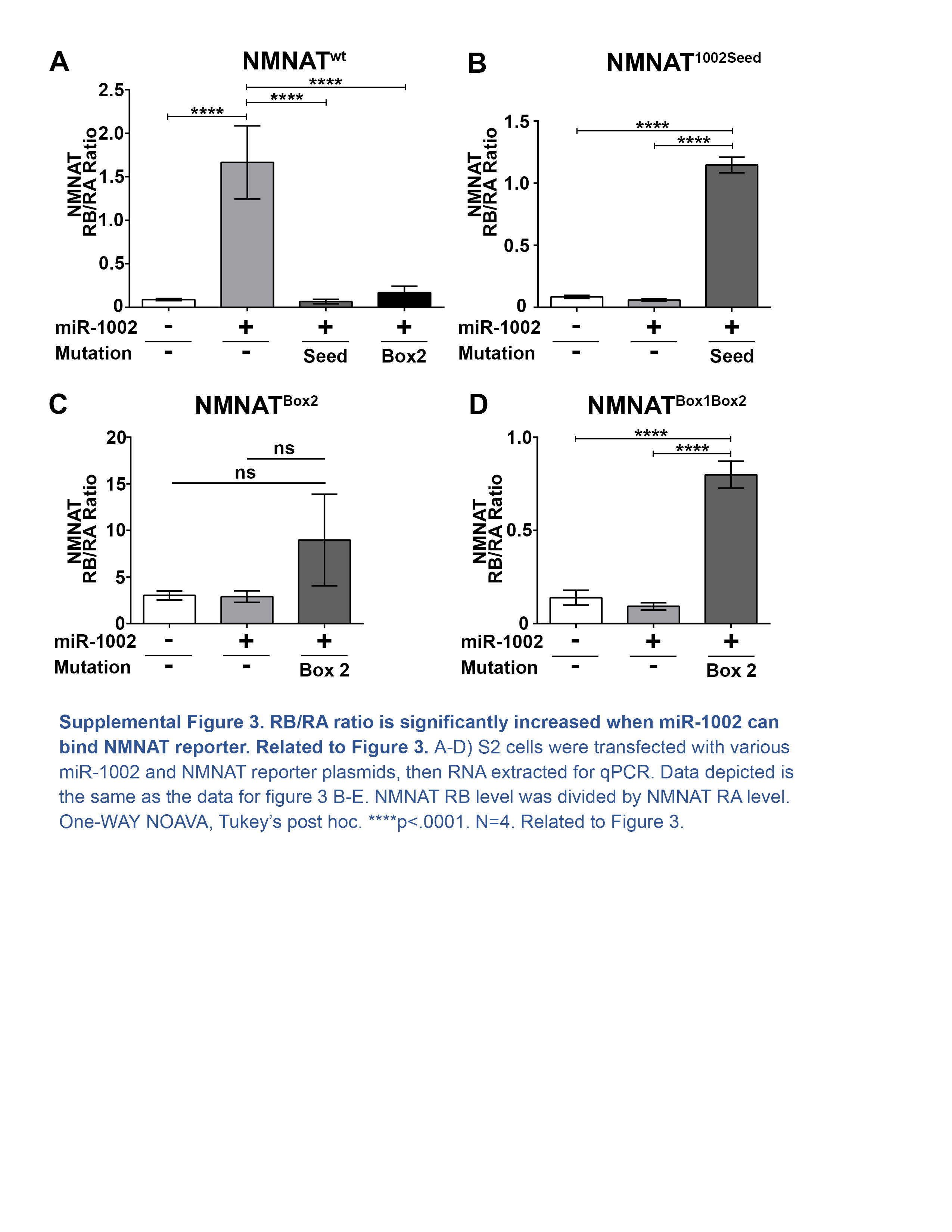
